# Predicting suitability of regions for Zika outbreaks with zero-inflated models trained using climate data

**DOI:** 10.1101/771279

**Authors:** Samantha Roth, Miranda Teboh-Ewungkem, Ming Li

## Abstract

In recent years, Zika spread through the Americas. This virus has been linked to Guillain-Barré syndrome, which can lead to paralysis, and microcephaly, a severe birth defect. Zika is primarily transmitted by *Aedes (Ae.) aegypti*, a mosquito whose geographic range has expanded and is anticipated to continue shifting as the climate changes.

We used statistical models to predict regional suitability for autochthonous Zika transmission using climatic variables. By suitability for Zika, we mean the potential for an outbreak to occur based on the climate’s habitability for *Ae. aegypti*. We trained zero-inflated Poisson (ZIP) and zero-inflated negative binomial (ZINB) regression models to predict Zika outbreak suitability using 20 subsets of climate variables for 102 regions. Variable subsets were selected for the final models based on importance to *Ae. aegypti* survival and their performance in aiding prediction of Zika-suitable regions. We determined the two best models to both be ZINB models. The best model’s regressors were winter mean temperature, yearly minimum temperature, and population, and the second-best model’s regressors were winter mean temperature and population.

These two models were then run on bias-corrected climate projections to predict future climate suitability for Zika, and they generated reasonable predictions. The predictions find that most of the sampled regions are expected to become more suitable for Zika outbreaks. The regions with the greatest risk have increasingly mild winters and high human populations. These predictions are based on the most extreme scenario for climate change, which we are currently on track for.

**Author Summary:** In recent years, Zika spread through the Americas. This virus has been linked to Guillain-Barré syndrome, which can lead to paralysis, and microcephaly, a severe birth defect. Zika is primarily transmitted by *Aedes (Ae.) aegypti*, a mosquito whose geographic range has expanded and is anticipated to continue shifting as the climate changes. We used statistical models to predict regional suitability for locally-acquired Zika cases using climatic variables. By suitability for Zika, we mean the potential for an outbreak to occur based on the climate’s habitability for *Ae. aegypti*. We trained statistical models to predict Zika outbreak suitability using 20 subsets of climate variables for 102 regions. Variable subsets were selected for the final two models based on importance to *Ae. aegypti* survival and their performance in aiding prediction of Zika-suitable regions. These two models were then run on climate projections to predict future climate suitability for Zika, and they generated reasonable predictions. The predictions find that most of the sampled regions are expected to become more suitable for Zika outbreaks. The regions with the greatest risk have high human populations and increasingly mild winters.

## Introduction

Recently, Zika has spread through many South American, Central American, and Caribbean countries. There have been thousands of autochthonous Zika cases in US territories like Puerto Rico, the US Virgin Islands, and American Samoa. In the mainland US, there have been over two-hundred autochthonous cases in Florida and a few in Texas [1]. In this paper, any Zika cases mentioned are autochthonous, meaning that the infection occurred in the given region. The spread of Zika and the impact of climate variation on its spread should be of concern to government officials, health professionals, and citizens in the Americas. Although the symptoms are usually mild and short-lived, Zika has been shown to be associated with Guillain-Barré, a disease that causes severe weakness of the muscles, and can sometimes lead to paralysis [2]. In addition, pregnant women who contract Zika can transmit the virus to their fetus, which can cause severe birth defects, such as microcephaly [2]. Currently, there is no vaccination or cure for Zika available, making it especially important to avoid contact with its vectors [3].

Since relatively less is known about the effects of climate on the Zika virus itself, we seek to model this relationship through the effects of climate on the vectors that spread Zika [4]. *Ae.* mosquitoes (*Ae. aegypti* and *Ae. albopictus*) transmit the Zika virus [5]. Both *Ae.* mosquitoes take shelter in human habitations, but *Ae. albopictus* primarily bites outside [4]. *Ae. albopictus* mosquitoes can undergo a winter diapause for up to a year as eggs, making them more capable of surviving in temperate climates than *Ae. aegypti* [6]. *Ae. aegypti* larvae develop more quickly at higher temperatures [7]. Larvae develop fastest at 32°C (89.6°F), while significant mortality occurs at or below 14°C (57.2°F) and at or above 38°C (100.4°F) [8]. The lower and upper limits for development are 16°C (60.8°F) and 34°C (93.2°F), respectively [9].

Temperature also impacts the ability of *Ae. aegypti* mosquitoes to fly and feed. They can sustainably fly between 15°C (59°F) and 32°C (89.6°F) [10]. For a short flight, the lower temperature limit is 10°C (50°F), and the upper limit is 35°C (95°F) [10]. In addition, they feed more quickly between 26°C (78.8°F) and 35°C (95°F) than between 19°C (66.2°F) and 25°C (77°F), [11]. *Ae. aegypti* does not bite below 15°C (59°F) [11],[12],[13]. *Ae. aegypti* are most active at 28°C (82.4°F) [12], and the optimal temperature for transmission is 29°C (84.2°F) [14]. Below 10 °C (50°F), *Ae. aegypti* cannot move, and above 40°C (104°F), they die [9]. In Japan and Asia, the minimum mean temperatures in winter months for a region to be habitable for *Ae. aegypti* were suggested to be −2 and −5 °C [15],[16]. There is less research concerning the effects of temperature on the flight and feeding behaviors of *Ae. albopictus* [4].

For both *Ae. aegypti* and *Ae. albopictus*, temperature has been shown to be the most crucial predictor of the vectors’ geographic distribution [17]. *Ae.* mosquitoes already have geographical ranges covering much of the Eastern United States. In terms of habitability for *Ae. aegypti*, the entire southeastern United States is a very likely range, with much of the Southern Appalachian area being a likely range [18]. The geographical range of the *Ae. albopictus* along the Eastern portion of the US is much larger, with the very likely range stretching all the way from the tip of Florida to the middle of Pennsylvania. The potential geographical regions for these two species of mosquitoes, as determined by the Centers for Disease Control and Prevention (CDC), are provided (Fig 1):

**Fig 1.**
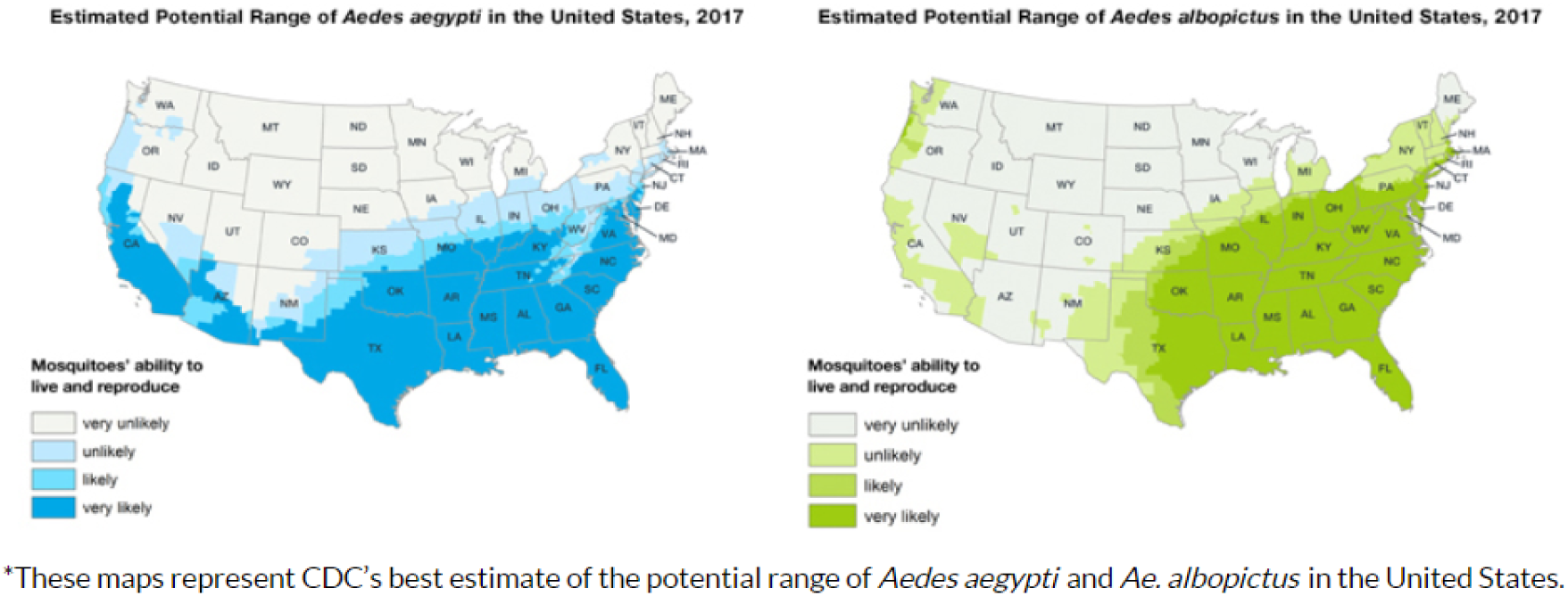
These figures are open-access content provided by [18].

Below (Fig 2), a 2015 estimate of the potential range of *Ae. aegypti* in the Americas is shown with permission from the authors.

**Fig 2.**
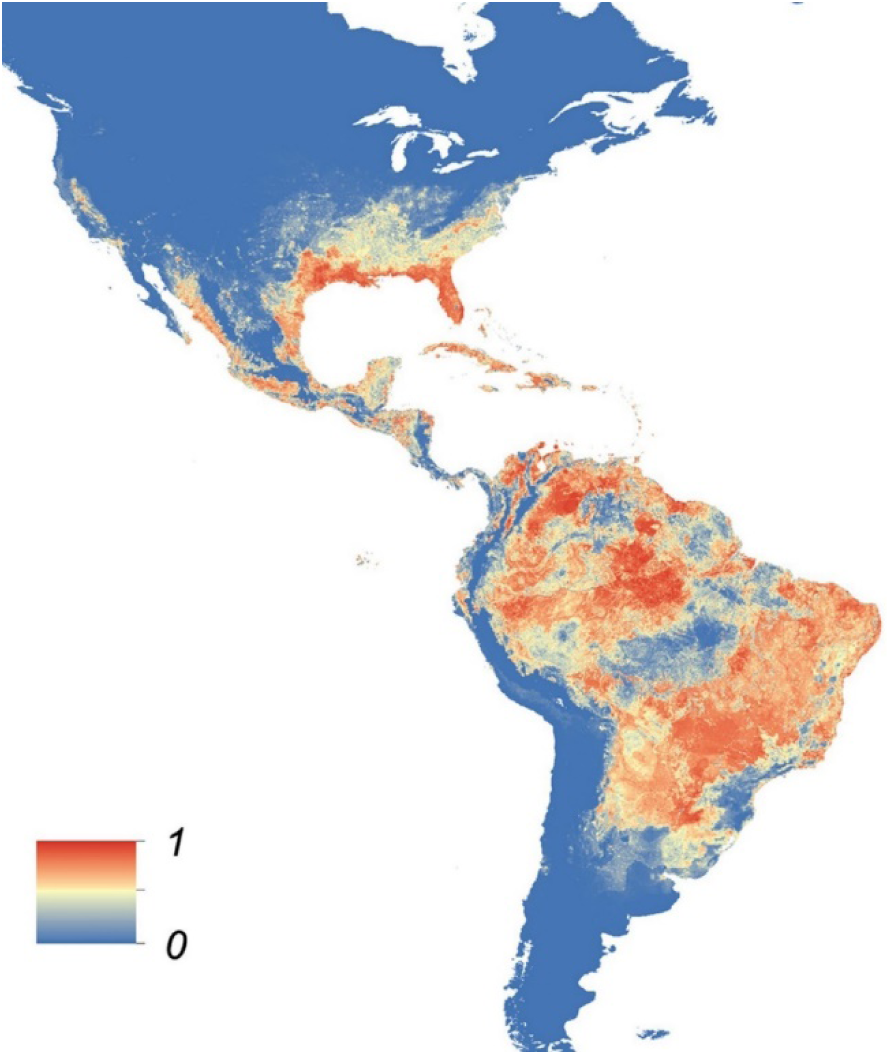
The colors are scaled from blue to red to indicate the probability of *Ae. aegypti* occurrence, where pure blue indicates zero probability, and pure red indicates a probability of one [17].

Recently, there were reports of *Ae. aegypti* in Santiago, Chile, a region that had previously eradicated this type of mosquito and had not expected to see its return [19]. Also, this map does not appear to reflect that *Ae. aegypti* have been detected in Las Vegas [20]. A 2018 map of the potential present distribution of *Ae. aegypti* [21] indicates that areas surrounding Portland, Oregon and Seattle, Washington would be suitable, but does not reflect the suitability of Las Vegas, Nevada [20] or Phoenix, Arizona [22]. The CDC’s map (Fig 1) also places all of Nevada in the “unlikely” or “very unlikely” ranges. These maps’ lack of including all regions where *Ae. aegypti* can survive indicate that they may need to be updated.

Currently, *Ae. aegypti* has trouble establishing populations in many temperate regions because of their colder winters; their eggs have a high mortality rate upon contacting frost [6]. In addition, there has been a decline in *Ae. aegypti* mosquitoes associated with the invasion of the *Ae. albopictus* mosquitoes [23]. *Ae. aegypti* mosquitoes prefer feeding on humans to other mammals, and feed numerous times during the gonotrophic cycle, increasing the risk of transmitting disease from host to host [6]. *Ae. albopictus* mosquitoes also prefer human blood, but they are opportunistic feeders and will feed preferentially on whatever mammals are available [24], whereas *Ae. aegypti* seek out human hosts even in the presence of other hosts [4],[6]. *Ae. albopictus* are less competent vectors and are therefore less likely to spread Zika than *Ae. aegypti* mosquitoes [24]. *Ae. aegypti* and *Ae. albopictus* are the primary vectors capable of transmitting Zika [3], [25].

Attributed to both climate change and El Niño, Latin America was hotter than average in 2015. There were high levels of precipitation in Southern Brazil and Uruguay in the winter leading into 2015 [26]. Rain leads to standing bodies of water in the form of puddles and water accumulated in other outdoors containers. These bodies of water serve as breeding grounds for *Ae. aegypti* mosquitoes [26]. However, too heavy of rainfall events can wash away larvae from breeding grounds, negatively affecting mosquito population size [27]. Also, artificial containers of water provide breeding sites [28], enabling mosquitoes to thrive during droughts [6],[29]. This is evinced by the start of the Zika outbreak of 2015 occurring in northeastern Brazil during a drought [26]. Results concerning the importance of precipitation to *Ae. aegypti* mosquitoes have been mixed, and heat was likely a more important factor in the Brazilian Zika epidemic. In warmer temperatures, viruses develop faster in vectors, inflating the proportion of infected mosquitoes [30]. There also tend to be more people outside and people with exposed skin to be easily infected [26]. It follows that maximal Zika transmission has been found to occur between 26 and 29°C [31].

Over the past few decades, *Ae. aegypti* has expanded its geographical range [6],[32]. Their distribution is expected to continue shifting as the climate shifts, putting new people at risk of Zika [6],[32],[33]. Since Zika is closely related to the dengue virus, and the two are both spread by *Ae. aegypti*, research relating climate to the spread of the dengue virus can also be applied to Zika [26],[34]. Temperature increases may expand *Ae. aegypti*’s range, increasing the range of the virus, but some areas may get too warm, leading to a decrease in *Ae. aegypti* and a decrease in virus transmission in some areas [26]. Therefore, research concerning climate change and Zika vectors is crucial for understanding how Zika risk in different regions may evolve in the coming decades.

Researchers have used a variety of methods to model the spread of Zika in different regions. In 2017, Zhang et al. used temperature, vector density, socioeconomic factors, human mobility, and demographic data in a “data-driven global stochastic epidemic model” to predict the spread of Zika in the Americas [35]. In 2018, Lo and Park used *Ae. aegypti* density, human density, and temperature data to perform a topological analysis to model the spread of Zika in Brazil [36]. Also in 2018, Huber, Caldwell, and Mordecai used a dynamic disease transmission model to examine the impact of different aspects of temperature: seasonal variation, seasonal mean, and value at epidemic onset on disease dynamics in twenty cities [14]. The New York City (NYC) Department of Health and Mental Hygiene applied a zero-inflated Poisson model to estimate which census tracts in NYC have the highest risk for Zika being imported by humans [37]. Other researchers have used SEIR model variations, time series models, and stochastic models for Zika-related research [36]. For other mosquito-related research, zero-inflated models have been used to model the relationship between weather and dengue incidences in China [38], mosquito egg abundance in Eastern Europe [39], *Ae. albopictus* females and egg abundance in France [40], and to analyze the density of adult malaria mosquitoes in Kenya [41]. In addition, the relationship between water volume changes in containers and *Ae. aegypti* pupal abundance has been explored in Vietnam [42]. Our method is unique in its relatively simple yet useful approach in characterizing a regions suitability for Zika outbreaks using only historical climate, population, and Zika data.

The magnitude of the climate variations we will experience depend on our efforts to reduce the amount of greenhouse gases (GHGs) being emitted. We can do a lot more to lower our emissions and not continue on the path aligning with the “business as usual” scenario, which accompanies the most extreme projections for climate change. In this manuscript, we explore the potential for Zika’s geographical reach to expand or shift with changes in climate by comparing Zika case predictions of statistical models trained using past climate data and future climate projections. Where by training a model, we mean providing an algorithm and data for the model to learn from. The overall goal of this research was to develop a statistical regression model that can identify regions capable of experiencing a Zika outbreak and to use this model to predict what regions may be at an increased risk for Zika outbreaks as the climate changes. As desired, we achieved this goal.

### Methodology

To quantify how climate variations may impact the potential spread of Zika outbreaks in the Americas, we sought out to build statistical models in R that could accurately identify regions that have already reported at least one Zika case and regions that presently have the potential for Zika infections to occur. Such models were created with the goal of predicting what areas may be impacted by Zika in the near future (2020-2045) given the variations in climate. To start, we constructed statistical models using various subsets of climate variables relevant to the survival of *Ae. aegypti*. The regression models we selected to compare were zero-inflated Poisson and zero-inflated negative binomial.

To gain a basis for understanding the zero-inflated Poisson regression model, we first consider the Poisson regression model. This is the simplest model in the family of generalized linear models (GLMs) [43]. The dispersion parameter is 1; the canonical link is *g*(*μ*_*i*_) = log (*μ*_*i*_), and the variance is *μ*_*i*_ [43]. Mathematically, the predictions from a Poisson model take the following form when there are *m* explanatory variables: log (*μ*_*i*_) = *β*_0_ + *β*_1_*x*_1_ + *β*_2_*x*_2_ + … + *β*_*m*_*x*_*m*_, where log (*μ*_*i*_) = log (*E*(*y*_*i*_)) is the natural log of the expected value of the response variable *y*_*i*_, and the *x*_*j*_s are the explanatory variables [44]. This is equivalent to: *μ*_*i*_ = exp (*β*_0_) exp (*β*_1_*x*_1_) exp (*β*_2_*x*_2_)…exp (*β*_*m*_*x*_*m*_), where exp (*β*_0_) is the effect on *μ*_*i*_ when all the *x*_*j*_s are zero, and exp (*β*_*j*_) is the multiplicative effect that every unit increase in *x*_*j*_ has on *μ*_*i*_ with all the other *x*_*k*_s held constant [44],[45]. Concerning the values of the *β*_*j*_s, if *β*_*j*_ = 0, then there is no association between *y* and *x*_*j*_; if *β*_*j*_ > 0, then *μ*_*i*_ increases when *x*_*j*_ increases, and if *β*_*j*_ < 0 then *μ*_*i*_ decreases when *x*_*j*_ increases [44].

A potential issue with Poisson models is overdispersion, which occurs when the variance of the data differs from the variance of a Poisson random variable [44],[45]. If overdispersion occurs, an adjustment can be made [44], or a quasi-Poisson [46] or negative binomial model could be used instead [44],[45]. If the overdispersion is due to a greater number of zeros than would occur in Poisson distribution, a zero-inflated Poisson (ZIP) or a zero-inflated negative binomial (ZINB) model should be considered [43],[45].

The next model we introduce is the count portion of the ZINB model: the negative binomial regression model. The probability density function for the negative binomial distribution takes the form: 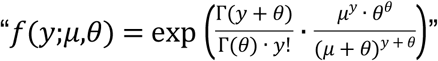 [43]. In this equation, *μ* is the mean; *θ* is the shape parameter, and Γ(·) is the gamma function [43]. The shape parameter can either be specified or estimated from the data. The negative binomial model has the same dispersion parameter as the Poisson model, but rather than being set equal to the mean, the variance is governed by the function: 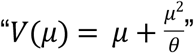 [43]. The negative binomial model for expected count has the same form as that of the Poisson model but accounts for over-dispersion using an additional parameter [44],[47].

The ZIP and ZINB regression models are useful when too many zeros are observed than would naturally occur in a Poisson or negative binomial distribution [45], [47]. This was the case with our dependent variable, Zika cases. A zero-inflated model is defined to be a “two-component mixture model combining a point mass (logistic regression model) with a count distribution”, which was either Poisson or negative binomial as in our case [43]. In the S1 Text, we summarize technical information on zero-inflated models provided by Zeileis, Kleiber, and Jackman [43] and Dr. Diane Lambert. If there are excess zeros, and there is still over dispersion when a ZIP model is used (i.e., over dispersion not from the excess zeros), a ZINB model may be a better fit [48]. As reflected through the models’ performances, this was the case with our data as well.

Next, we acquired all observational and projected climate data through Climate Change, Agriculture, and Food Security (CCAFS) Climate Portal [49]. Originally, each region that reported Zika cases was going to be represented by the yearly temperature and precipitation data for the years 2015 to 2017, since those were the years that Zika data was available from the Pan-American Health Organization (PAHO) [50]. However, for some of the areas outside of the United States, the necessary weather data was not available for those years from reliable resources. For the sake of consistency and reliability, we chose to use historical data from 1980-2005 from CCAFS, since it was available for all the necessary regions in North and South America. Some Caribbean islands that had Zika cases were not available through this source.

Originally, we were going to use data for each US state and territory and for each country outside the US. However, only looking at the number of Zika cases and the climate at the country level for all nations other than the US and at the state level for some US states would leave out significant climate variation across some nations. So, we chose to further break down the following nations and US states: Brazil, Argentina, Bolivia, Paraguay, Texas, Mexico, Canada, California and Florida. Since the highest level of detail in reporting Zika cases was at the district/state level in the selected South American countries, we chose a point in each state or district to represent the climate there. Almost all of these points represented a city in the state/district, with the exception of La Paz, Bolivia. Bolivia ranges from low-lying zones with warmer temperatures to colder mountainous zones, and the district of La Paz contains both of these [51],[52]. Since the largest city in La Paz, also named La Paz, is in the mountainous region deemed unlikely to be inhabited by *Ae. aegypti*, we chose a point in a lower-lying, warmer part of the district of La Paz instead [51]. For the selected cities, we selected a longitude and latitude point in the center of the dot representing the city and downloaded the historical climate data and the bias-corrected past future projections. We chose to use cities as representative points since *Ae. aegypti* tend to live near people and bite inside buildings.

CCAFS obtained historical climate data from Agricultural Climate Forecast System Reanalysis (AgCFSR), which combines “daily resolution data from retrospective analyses (…the Climate Forecast System Reanalysis, CFSR) with in situ and remotely-sensed observational datasets” [53]. The uncorrected historical and future climate projections were from the Beijing Climate Center Climate System Model 1.1 in the Coupled Model Intercomparison Project 5 (CMIP5). The CCAFS Climate Portal authors then bias-corrected these projections using quantile mapping [49].

The future of the climate is uncertain and depends on the emissions levels today and in the years to come. So, for the future climate projections, we selected the emissions scenario that we are currently on track for, Representative Concentrations Pathway 8.5 (RCP 8.5). This is the highest emissions scenario explored by the CMIP5 climate modelers, so the temperature and precipitation changes in the future scenarios are expected to be more extreme than in lower emissions scenarios (RCPs 2.6, 4.5, and 6). This scenario is also interesting to consider in terms of its potential impact on the habitable areas for *Ae. aegypti* mosquitoes, because currently they are limited from spreading as far north as *Ae. albopictus* mosquitoes have because their eggs cannot survive the cold winters. With these winters potentially becoming warmer under this emissions scenario, *Ae. aegypti* mosquitoes could potentially inhabit a wider range of areas. However, some regions that are warm enough for these mosquitoes could become too hot in the summer and experience a decline in *Ae. aegypti* populations, and in turn, Zika infections.

Since the CCAFS climate data was not available for the years in which the Zika outbreaks occurred, we compared the historical climate data to monthly weather data provided by Weather Underground from the years that Zika was reported in the Americas (2015-2017) [54]. This data was not used in the model because the weather reports for 2015-2017 were either filled with missing/bad data or entirely unavailable for some of the Zika-positive regions. For many of the Zika-positive regions, the weather reports contained a few unrealistic or missing observations. These “bad” observations were identified using judgment based on the reported temperatures from the rest of the month and the climate of the region. To replace unrealistic or missing values, we used “seasonally decomposed missing value imputation” [55]. In R, the function that performs seasonally decomposed missing value imputation is called *na.seadec* and the package containing this function is package *imputeTS* [55]. This function first eliminates the seasonal component, imputes missing values using linear interpolation, then reinstates the seasonal component [55]. After performing seasonally decomposed missing value interpolation, we calculated the statistics *winter.mean*, *summer.mean*, *year.min*, *year.mean*, *year.max*, *year.prec*, and *year.diff* for the years 2015-2017 for Zika-reporting regions. We then compared these statistics to those calculated from the 1980-2005 CCAFS climate data by computing the mean and median differences between them and the mean absolute differences as shown in Table 1.

**Table 1.** The calculations behind this summary data were performed using Excel. For each climate statistic, Mean Absolute Difference is defined as the mean of the absolute values of the 2015-2017 means minus the 1980-2005 means. Mean Difference is defined as the mean of the 2015-2017 means minus the 1980-2005 means. Median Difference is defined as the median of the 2015-2017 means minus the 1980-2005 means.

| Climate Statistic | Mean Absolute Difference | Mean Difference | Median Difference |
| --- | --- | --- | --- |
| winter.mean | 1.4267 °C | 1.1546 °C | 1.0442 °C |
| summer.mean | 1.2720 °C | 0.9227 °C | 0.8109 °C |
| year.min | 2.8734 °C | 1.5114 °C | 1.1963 °C |
| year.mean | 1.2389 °C | 0.9697 °C | 0.7782 °C |
| year.mean15to32 | 0 | 0 | 0 |
| year.max | 3.0824 °C | -2.1741 °C | -2.2177 °C |
| year.prec | 971.4402 mm | -619.5307 mm | -472.663 mm |
| year.diff | 1.4448 °C | -0.9396 °C | -0.8096 °C |

We chose the climate variables to be used in the models based on their importance to *Ae. aegypti*. Since temperature is the most important factor in determining *Ae. aegypti*’s geographical range, most of the variables were chosen to describe the central tendency and spread of temperature throughout the year. Initially, we were going to include precipitation since it plays a role in breeding ground availability for *Ae. aegypti*. However, there were major discrepancies in precipitation between the years the Zika outbreak occurred and the years in the historical period used to train the models. So, we eliminated this variable. We included the mean difference between daily maximum and minimum temperature since large temperature changes in 24 hours negatively impacts female reproduction [56]. The climate variables considered were: mean yearly minimum temperature (*year.min*), mean yearly mean temperature (*year.mean*), mean yearly maximum temperature (*year.max*), an indicator for *year.mean* being within the range in which *Ae. aegypti* can sustainably fly (*year.mean15to32*), the mean difference between the daily maximum, and daily minimum temperatures (*year.diff*), the mean yearly winter mean temperature (*winter.mean*), and the mean yearly summer mean temperature (*summer.mean*).

To obtain the average yearly minimum (mean) (maximum) temperature, we took the minimum (mean) (maximum) of all the daily minimum temperatures for all the days for each year. Then we averaged the yearly minimums (means) (maximums) for minimum (mean) (maximum) daily temperature over the 21-year period. We calculated *winter.mean* and *summer.mean* by taking the mean of the mean temperatures for December, January, and February, the taking the mean of the mean temperatures for June, July, and August, and then assigning one of these means to winter and one to summer depending on whether the region of interest is above or below the equator.

Counts of Zika cases per country were obtained from PAHO; cases in US states were obtained from the CDC, and cases in states/districts of non-US countries that were further broken down this way were obtained from their respective health surveillance websites and PAHO [57],[58],[59],[60]. First, we gathered the most recent Zika data made available on the PAHO website, which was dated 1/4/2018 [50]. For regions that were divided into smaller regions, this data set was used to scale the more specific geographical distribution data. Scaling was necessary because the geographical distribution data was found in other sources reported at different time periods, leading them to have different reported numbers of Zika cases for each region of interest than was reported in the final PAHO report. These sources include earlier PAHO reports for individual South American countries that were dated before the latest report was released and the country/states’ websites when they were available. Paraguay’s website where PAHO obtained its data was no longer available. So, for selected departments (Amambay and the Metropolitan Area of Asuncion), we took the total number of cases in 2016, occurring in Amambay, the Metropolitan Area of Asuncion, Alto Parana, and Paraguari, and scaled them by the areas’ proportion of the population where the cases were observed to estimate numbers of cases for specific departments [60]. All of the Zika and climate data was recorded in Excel spreadsheets, which are available in the supplementary information. Each row of the spreadsheet has data for the point representing the specified department, state, or city in the specified region.

Next, we performed the exploratory analysis by creating histograms for all of the independent variables and the dependent variable, *Cases* (number of Zika cases up to 2018). The histogram of *Cases* revealed a much larger number of zeros than would have been observed in the Poisson or negative binomial distribution, so we decided that a zero-inflated model would likely perform best. We also noticed that the standard deviation of *Cases* was much larger than the mean, indicating over dispersion possibly due to the excess zeros. So, we decided to train both the zero-inflated negative binomial and Poisson models using different subsets of the climate statistics as regressors and compare their performances in predicting Zika outbreak suitability.

Each of 40 models (20 ZIP and 20 ZINB) created was trained on the historical climate and Zika data using all rows minus one, then run on the row left out of the training set to predict the number of Zika cases for that region. For each model, this process repeated 101 more times so we could get the model’s prediction for each region without training the model using that region. In terms of regressors used in these models, we started with the simplest subset first: only one regressor. The variable chosen was *year.min*, since susceptibility to cold is a limiting factor in the *Ae. aegypti’s* distribution. Next, we tested the performance of a simple model using another measure of coldness: *winter.mean*. Since there were over 1,000 potential models, we did not test all possible model; not all combinations of regressors would have been logical to test anyway. Instead, we chose 20 subsets of climate variables; each subset would be tested using a ZIP and a ZINB model. The model performances were evaluated based on soundness of model coefficients (i.e., coefficients that should be positive were positive and coefficients that should be negative were negative), ability to predict at least one Zika case for all regions that reported Zika cases, and for regions that did not yet report Zika, only predicting one case or more for regions that presently could experience a Zika outbreak. We define Zika-suitable regions as those currently inhabited by or determined to be capable of being inhabited by *Ae. aegypti* by at least two out of three of the potential distributions discussed in the introduction. In the potential distribution provided by the CDC, we considered an area in the “likely” or “very likely” range to be capable of being inhabited by *Ae. aegypti*. The presence of *Ae. aegypti* is correlated to Zika outbreaks if there are individuals in that region with Zika. Since we used count models, we considered a prediction of at least one Zika case to indicate that the model considers a region suitable for Zika.

For the regions that reported Zika, we compared each model’s predictions to the observed cases by calculating the mean of the absolute values of the residuals, then we scaled these values by dividing them by the number of cases observed to get the mean scaled residual. The mean scaled residual values were considered secondarily to the other methods of evaluating model performance. We determined that the zero-inflated negative binomial models consistently outperformed the zero-inflated Poisson models. The final models were two ZINB models (ZINB1 and ZINB2). The best performing model (ZINB1) used *winter.mean* and *year.mean* in the zero-inflation model and the same variables plus *Pop.100.000* in the count model. The second best performing model (ZINB2) used *winter.mean* only in the zero-inflation model and *winter.mean* plus *Pop.100.000* in the count model. These models were then run on future climate data to estimate where Zika outbreaks may pose a threat in the years 2020-2045.

## Results

### 1980-2005 vs 2015-2017 Climate Comparison

We compared the statistics calculated with the 1980-2005 data to those calculated with the 2015-2017 data. The 2015-2017 climate statistics can be found in the appendix.

On average, both the winters and summers of 2015-2017 were near 1 °C warmer for Zika-positive regions than they were in 1980-2005. The increase in *winter.mean* reflects an increase in Zika suitability between these two time frames. The average yearly minimum and yearly mean temperatures were also higher in general for 2015-2017 than they were for 1980-2005, also reflecting an increase in Zika suitability. The average yearly maximum temperature was lower in 2015-2017 than for 1980-2005. If the yearly maximum was near or above 32 °C, the yearly maximum decreasing from 1980-2005 to 2015-2017 could increase the Zika suitability of regions overall. The average difference between the daily minimum temperature decreased from 1980-2005 to 2015-2017, likely reflecting an increase in Zika suitability. As previously stated in the introduction, the relationship between precipitation and habitability for the mosquitoes that transmit Zika is complex, and the two time frames have average yearly precipitations very different from each other, so this variable was eliminated.

### Model Analysis-Present Predictions

Figs 3-8: A, C were created using the maps package in R. Longitudinal and latitudinal data were added using the *maps* and *ggplot2* packages in R. See the relevant code in the S3 Text for additional information.

#### Zero-Inflated Negative Binomial Regression Model 1 (ZINB1)

**Table 2.**
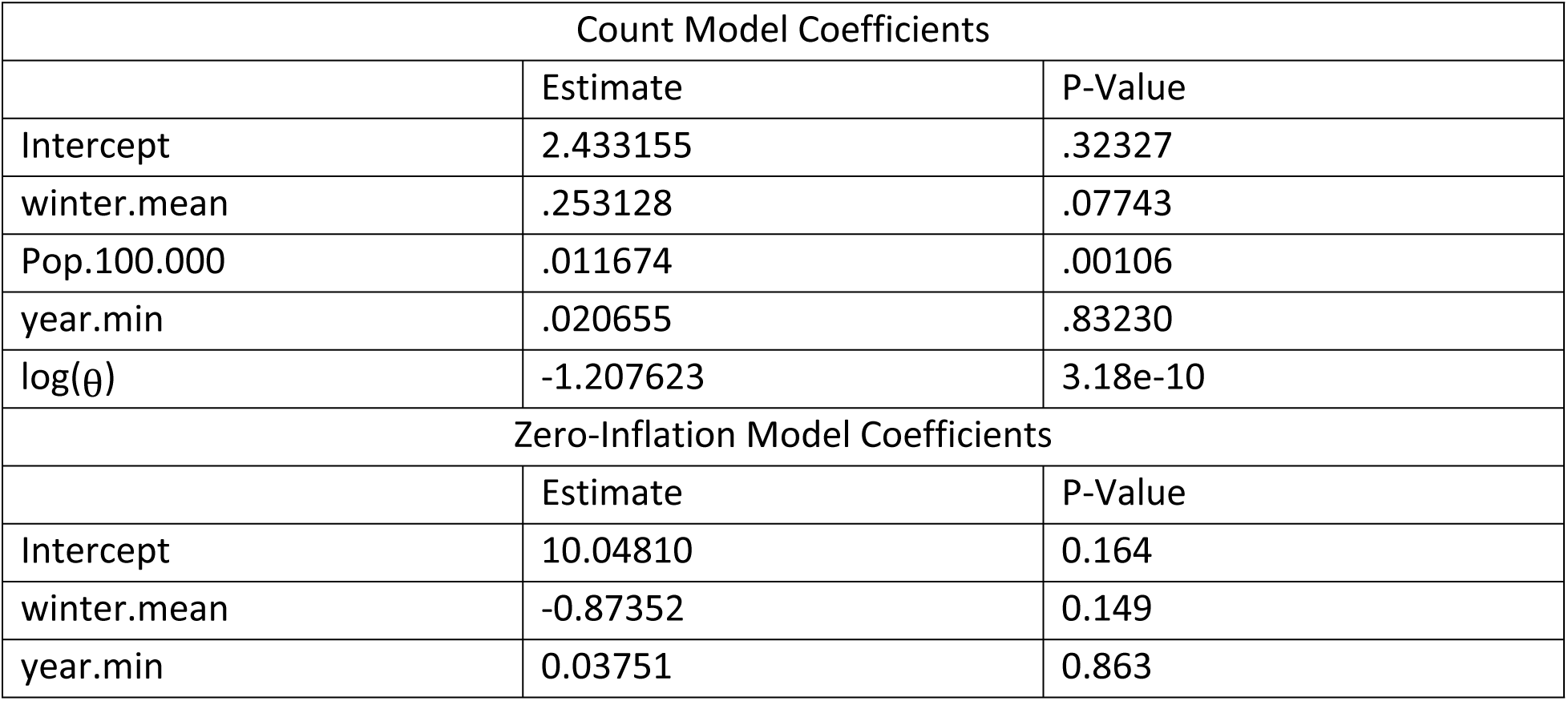
The coefficients and p-values in the table were produced using the zeroinfl() function in RStudio.

Starting with the output of the count model, log(_***θ***_) estimates the log of the inverse of *α*, the dispersion parameter, so the estimate of *α* is approximately 3.3455. The other coefficients in the count model represent the expected change in the log of the number of cases when the other variables are held constant [61]. So, holding the other regressors constant, a 1 °C increase in *winter.mean* changes the log of the predicted cases by .253128, an increase in population of 100,000 increases the log of the predicted cases by .011674, and a 1 °C increase in *year.min* increases the log of the predicted cases by .020655. Turning to the zero-inflation model, the other regressor held constant, an increase of 1 °C of *winter.mean* changes the log-odds of a region observing at least one case by -.87352, and an increase of 1 °C of *year.min* changes the log-odds of a region observing at least one case by .03751. Since all the coefficients in the count model are positive, the model’s coefficients are satisfactory.

Concerning the model’s predictions for Zika-free regions, the median is .007292707; the mean is 169.0705, and the standard deviation is 986.7907. Fig 3A below shows the geographic distribution of ZINB1’s predictions for Zika-free regions that were at least one. The Zika-free regions with the highest predictions were Los Angeles, California; Orlando, Florida; Louisiana; Uruguay; Cordoba, Argentina; Houston, Texas; Arizona; San Francisco, California; Nevada, and Mississippi. Fig 3B shows the distribution of predictions for Zika-free regions. For the Zika-positive regions, the model predicted a median of 7177.149, a mean of 28795.99, and a standard deviation of 92643.17. Fig 3C below shows the geographic distribution of ZINB1’s predictions for Zika-positive regions. All predictions for these regions were at least one. The Zika-positive regions with the highest predictions were Peru; Venezuela; Miami, Florida; Haiti; Nicaragua; Brownsville, Texas; the Dominican Republic; Guyana; Manaus, Brazil; and Panama. Fig 3D shows the distribution of predictions for Zika-positive regions. The model’s performance in predicting the number of cases reported in each Zika-positive region was measured using scaled residuals. For Zika-positive regions, the scaled residuals had a mean of 205.1306, a median of 4.497625, and a standard deviation of 31923.67.

**Fig 3.**
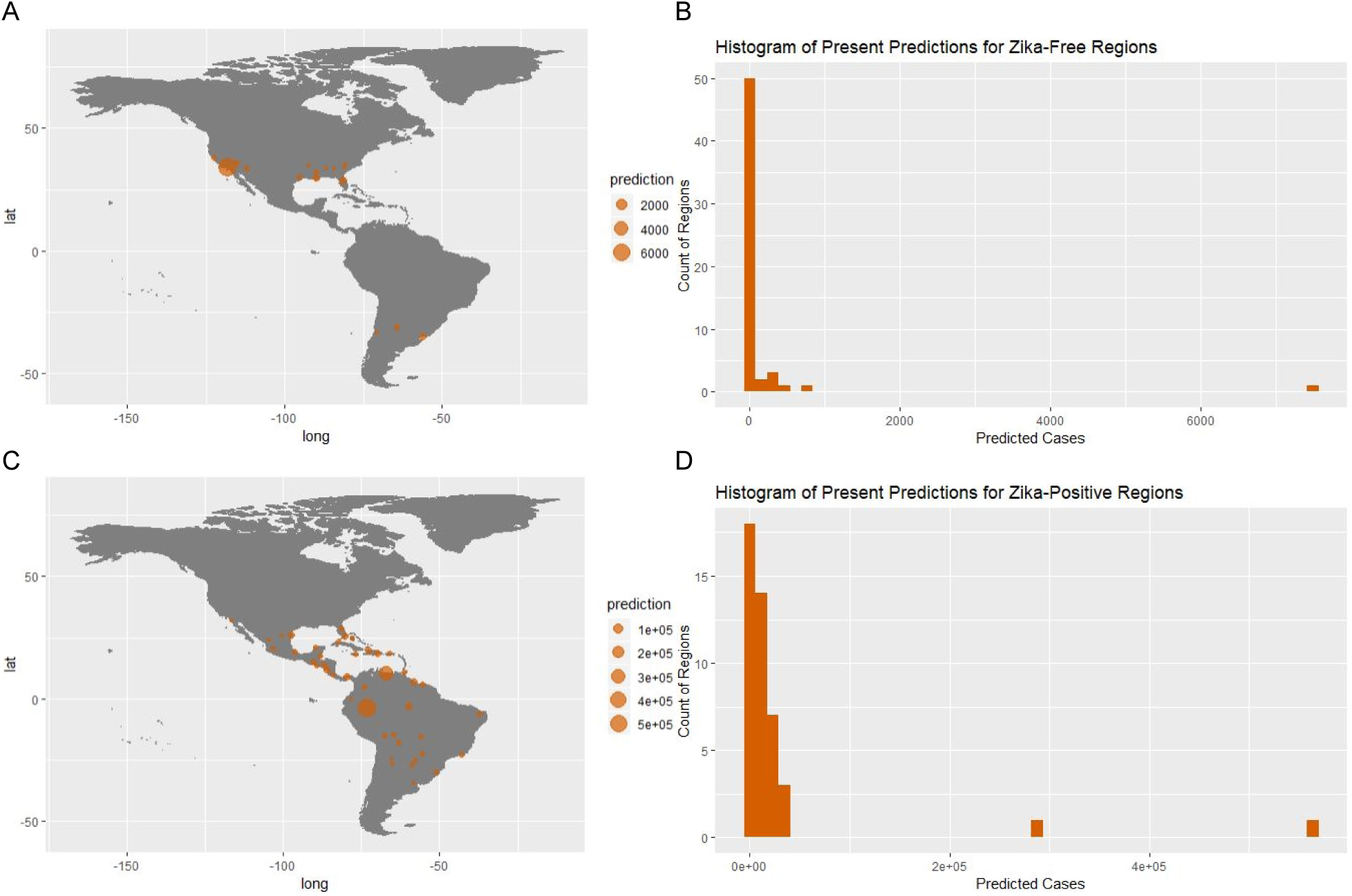
Fig A shows ZINB1’s present predictions for the Zika-free regions and was produced using the packages maps and ggplot2 in R. Fig B shows the distribution of predicted Zika cases for Zika-free regions and was produced using ggplot2 in R. Fig C shows ZINB1’s present predictions for the Zika-positive regions and was produced in the same manner as Fig A. Fig D shows the distribution of predicted Zika cases for Zika-positive regions and was produced in the same manner as Fig B.

#### Zero-Inflated Negative Binomial Regression Model 2 (ZINB2)

**Table 3.** The coefficients and p-values in the table were produced using the *pscl* package in RStudio.

| Count Model Coefficients |  |  |
| --- | --- | --- |
|  | Estimate | P-Value |
| Intercept | 2.068006 | 0.189355 |
| winter.mean | 0.278441 | 5.30e-05 |
| Pop.100.000 | 0.011834 | 0.000532 |
| $\log(\theta)$ | -1.197894 | 2.66e-10 |
| Zero-Inflation Model Coefficient |  |  |
|  | Estimate | P-Value |
| Intercept | 8.9361 | 0.033 |
| winter.mean | -0.7836 | 0.040 |

Beginning with the count model’s output, based on log(_***θ***_), the estimate of the dispersion parameter is about 3.3131. Holding the other regressors constant, a 1°C increase in *winter.mean* changes the log of the predicted cases by 0.278441, and an increase of 100,000 in populations increases the log of the predicted cases by 0.011834. In the zero-inflation model, an increase of 1°C of *winter.mean* changes the log-odds of a region observing at least one case by −0.7836. Again, since all the coefficients in the count model are positive, they are satisfactory.

For the Zika-free regions, the model predicted a median of 0.0102, a mean of 146.0512, and a standard deviation of 828.4233. Fig 4A shows the geographic distribution of ZINB2’s predictions of at least one for Zika-free regions. Fig 4B shows the distribution of predictions for Zika-free regions. The Zika-free regions with the highest predictions, i.e. predicted to have the most Zika-suitable climates, were: Los Angeles, California; Orlando, Florida; Louisiana; Uruguay; Cordoba, Argentina; Houston, Texas; Arizona; San Francisco, California; Las Vegas, Nevada; and Mississippi.

**Fig 4.**
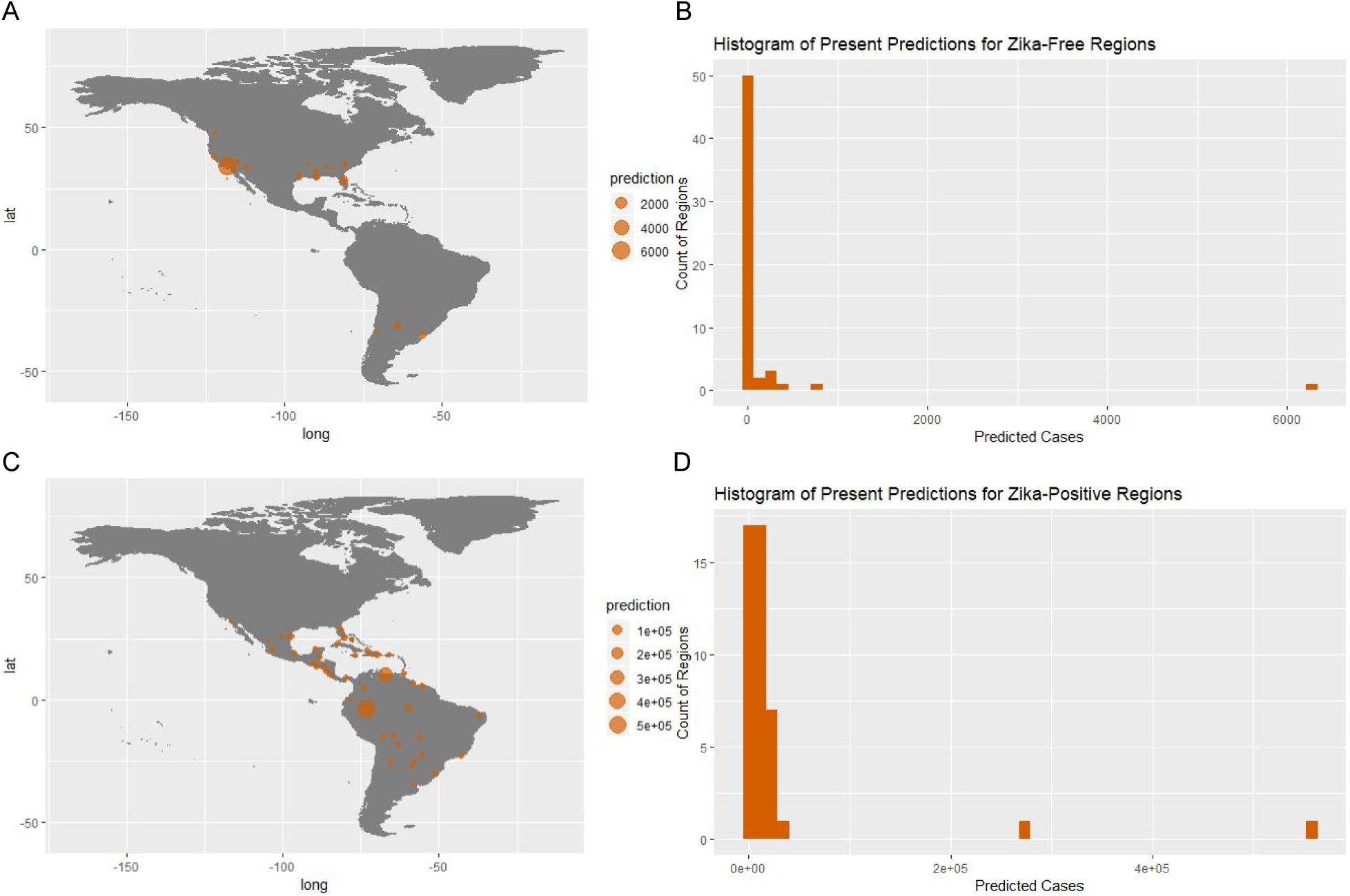
Fig A shows ZINB2’s present predictions for the Zika-free regions and was produced using the packages maps and ggplot2 in R. Fig B shows the distribution of predicted Zika cases for Zika-free regions and was produced using ggplot2 in R. Fig C shows ZINB2’s present predictions for the Zika-positive regions and was produced in the same manner as Fig A. Fig D shows the distribution of predicted Zika cases for Zika-positive regions and was produced in the same manner as Fig B.

For the Zika-positive regions, the model predicted a mean of 27885.65, a median of 8116.381, and a standard deviation of 91221.87. The Zika-positive regions predicted to have the most Zika-suitable climates were Peru; Venezuela; Miami, Florida; Nicaragua; Brownsville, Texas; Haiti; Dominican Republic; Manaus, Brazil; Panama; and Guyana. Fig 4C below shows the geographic distribution of ZINB2’s predictions for Zika-positive regions that were at least one. Fig 4D shows the distribution of predictions for Zika-positive regions. Concerning scaled residuals, the mean was 200.3012; the median was 4.252153, and the standard deviation was 781.1309.

### Model Analysis- Future Predictions (2020- 2045)

#### Zero-Inflated Negative Binomial Model 1 (ZINB1)

For the currently Zika-free regions, in the future the model predicted a mean of 375.9988; a median of 0.08198095, and a standard deviation of 2356.484. The histogram for future predicted cases is provided in Fig 5B. As shown in Fig 5A, the regions predicted to have the most suitable climate for Zika outbreaks (having the highest numbers of cases predicted), include Los Angeles, California; Louisiana; Orlando, Florida; San Francisco, California; Houston, Texas; Uruguay; Arizona; Cordoba, Argentina; Mississippi; and Nevada.

**Fig 5.**
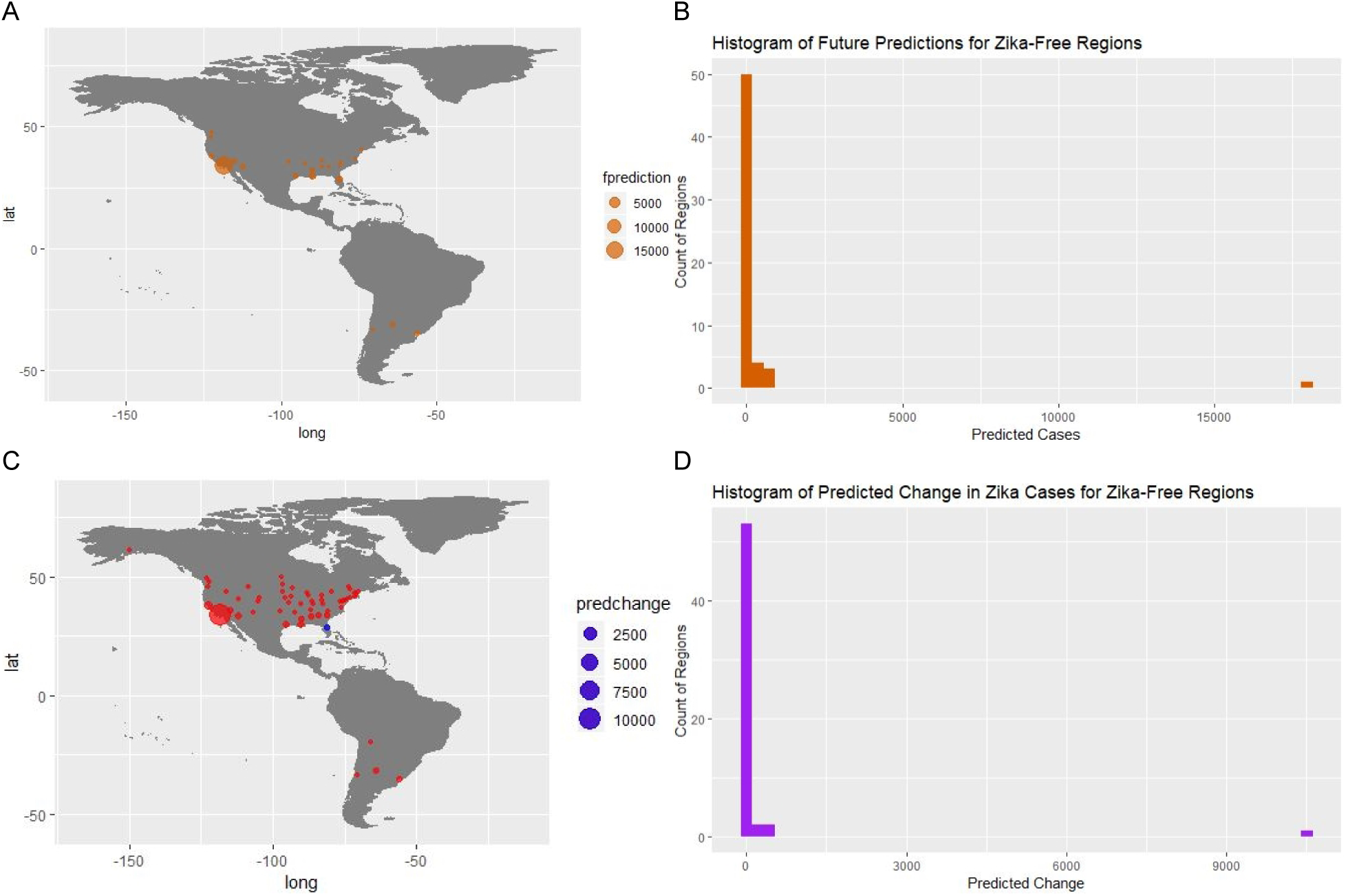
Fig A shows ZINB1’s future predictions for the Zika-free regions and was produced using the packages maps and ggplot2 in R. In the key, *fprediction* means future prediction. Fig B shows the distrbution of future predictions for Zika-free regions and was produced using ggplot2 in R. Fig C shows ZINB1’s change in predictions for the Zika-free regions and was produced in the same manner as Fig A. In Fig C, *predchange* means predicted change in cases, and the key shows the absolute value of the predicted change corresponding to the given dot size. On the map, a red dot represents a predicted increase, and a blue dot represents a predicted decrease. Fig D shows the distribution of the change in predictions for Zika-free regions and was produced in the same manner as Fig B.

Concerning the predicted changes in cases, the mean is 206.9283; the median is 0.06621878, and the standard deviation is 1374.127. The distribution of change in predictions is provided in Fig 5D. As shown in Fig 5C, the Zika-free regions predicted to see the greatest increases in Zika suitability include: Los Angeles, California; San Francisco, California; Louisiana; Houston, Texas; Arizona; Mississippi; Uruguay; Cordoba, Argentina; Nevada; and South Carolina. The only Zika-free region predicted to see a decrease in Zika suitability was Orlando, Florida. However, as previously stated, this region is still predicted to be quite suitable for Zika.

Shifting focus to the Zika-positive regions, ZINB1 predicts that on average, the regions that have already seen Zika will continue to be more Zika-friendly on average than the Zika-free regions. For the future predictions, the mean is 28795.99; the median is 7177.149, and the standard deviation is 92643.17. As shown in Fig 6A, the regions predicted to see the most Zika-suitable climates in the future period include Peru; Venezuela; Colombia; Haiti; Nicaragua; the Dominican Republic; Panama; Suriname; Miami, Florida; and Brownsville, Texas. As shown in Fig 6D, for changes in predictions, the mean is 5033.435, the median is 3157.645, and the standard deviation is 19710.42, indicating that these regions are expected to become more suitable for Zika outbreaks based on the climate projections considered. As shown in Fig 6C, the regions that have reported Zika that are expected to see the greatest increases in Zika outbreak suitability are Venezuela; Colombia; Haiti; the Dominican Republic; Nicaragua; Panama; Suriname; Puerto Rico; Jamaica; and Rio de Janeiro, Brazil. The only two presently Zika-positive regions predicted to experience a decrease in Zika outbreak suitability were Peru and Miami, Florida.

**Fig 6.**
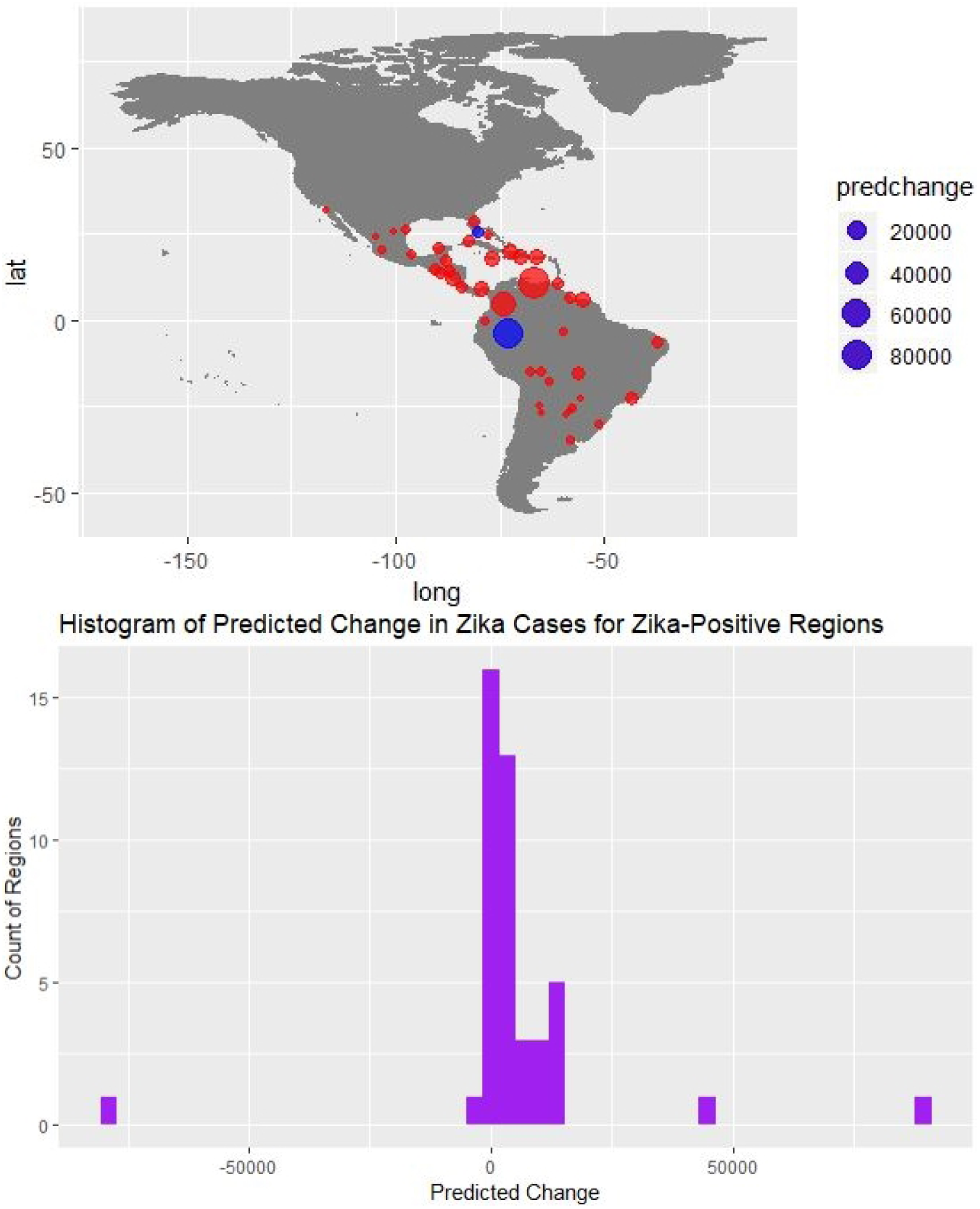
Fig A shows ZINB1’s future predictions for the Zika-positive regions and was produced using the packages maps and ggplot2 in R. In the key, *fprediction* means future prediction. Fig B shows the distribution of future predictions for Zika-positive regions and was produced using ggplot2 in R. Fig C shows ZINB1’s change in predictions for the Zika-positive regions and was produced in the same manner as Fig A. In Fig C, *predchange* means predicted change in cases, and the key shows the absolute value of the predicted change corresponding to the given dot size. On the map, a red dot represents a predicted increase, and a blue dot represents a predicted decrease. Fig D shows the distribution of predicted change in Zika cases for Zika-positive regions and was produced in the same way as Fig B.

#### Zero-Inflated Negative Binomial Model 2 (ZINB2)

For the predictions in Zika-free regions, the median is 0.0907; the mean is 387.0052, and the standard deviation is 2398.057, as shown in Fig 7B. As shown in Fig 7A, the regions predicted to be the most suitable for Zika in 2020-2045 are: Los Angeles, California; Orlando, Florida; Louisiana; Houston, Texas; San Francisco, California; Uruguay; Arizona; Cordoba, Argentina; Mississippi; and Nevada. For the Zika-free regions, the median predicted change in cases is 0.08406048; the mean is 240.954, and the standard deviation is 1573.279. As shown in Fig 7C, the Zika-free regions expected to see the greatest increases in Zika outbreak suitability are Los Angeles, California; Louisiana; San Francisco, California; Houston, Texas; Orlando, Florida; Arizona; Uruguay; Cordoba, Argentina; Mississippi, and Nevada.

**Fig 7.**
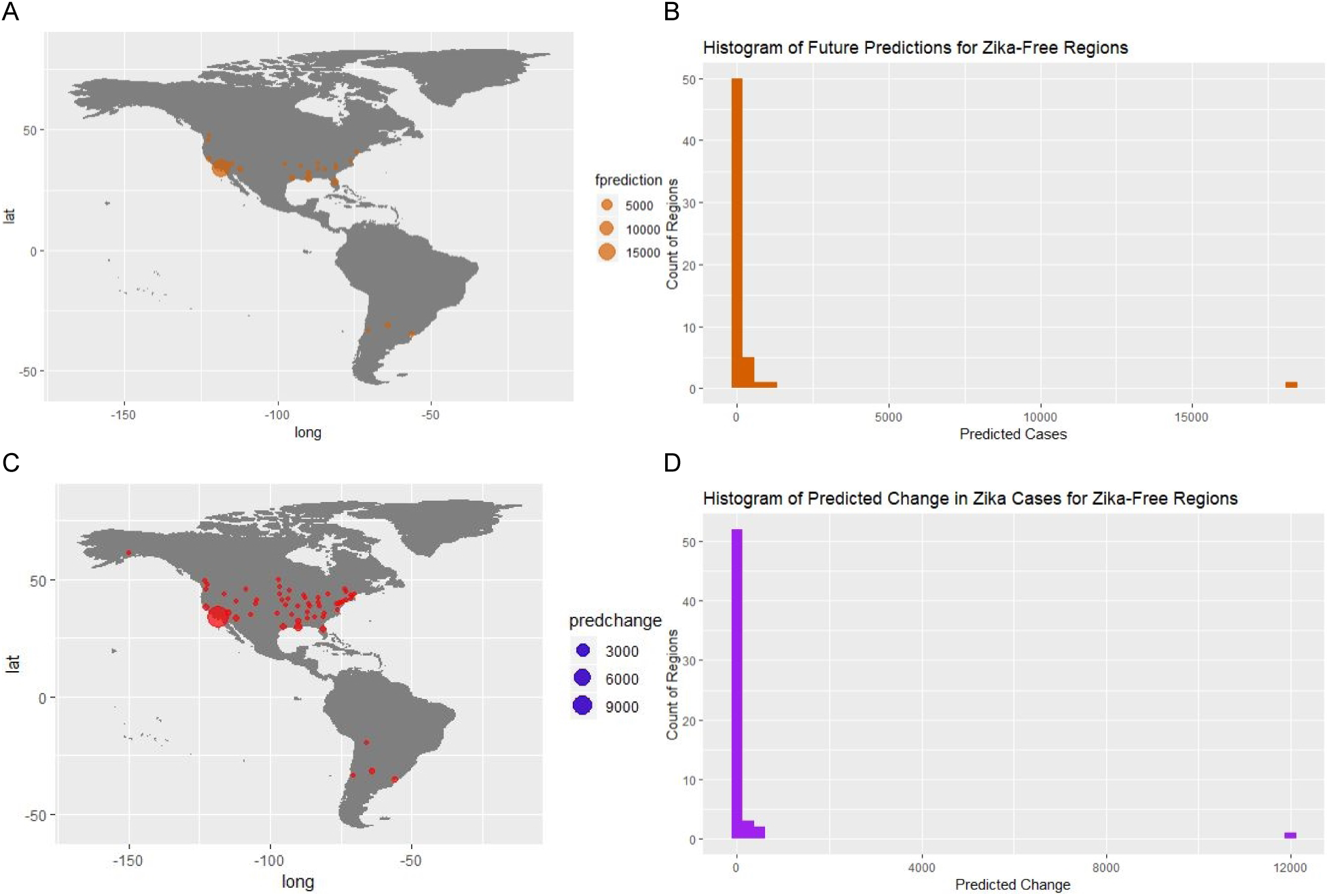
Fig A shows ZINB2’s future predictions for the Zika-free regions and was produced using the packages maps and ggplot2 in R. In the key, *fprediction* means future prediction. Fig B shows the distribution of future predictions for Zika-free regions and was produced using ggplot2 in R. Fig C shows ZINB2’s change in predictions for the Zika-free regions and was produced in the same manner as Fig A. In Fig C, *predchange* means predicted change in cases, and the key shows the absolute value of the predicted change corresponding to the given dot size. On the map, a red dot represents a predicted increase, and a blue dot represents a predicted decrease. Fig D shows the distribution of predicted change in Zika cases for Zika-free regions and was produced in the same way as Fig B.

As shown in Fig 8B, for the Zika-positive regions, the median prediction is 13502.49; the mean is 33799.84, and the standard deviation is 88408.86. As shown in Fig 8C, the currently Zika-positive regions predicted to be the most suitable for Zika in 2020-2045 are: Peru; Venezuela; Colombia; Nicaragua; Haiti; Miami, Florida; the Dominican Republic; Brownsville, Texas; Panama; and Suriname. For the Zika-positive regions, the median change in prediction is 4341.834; the mean is 5914.191, and the standard deviation is 17188.65. As shown in Fig 8D, ZINB2 predicts that all but one Zika-positive region will continue to become more Zika-friendly. As shown in Fig 8C, the Zika-positive regions expected to see the greatest increases in Zika outbreak suitability are Venezuela, Colombia, Haiti, Nicaragua, the Dominican Republic, Suriname, Panama, Jamaica, Rio de Janeiro, Brazil; Brownsville, Texas; and Puerto Rico.

**Fig 8.**
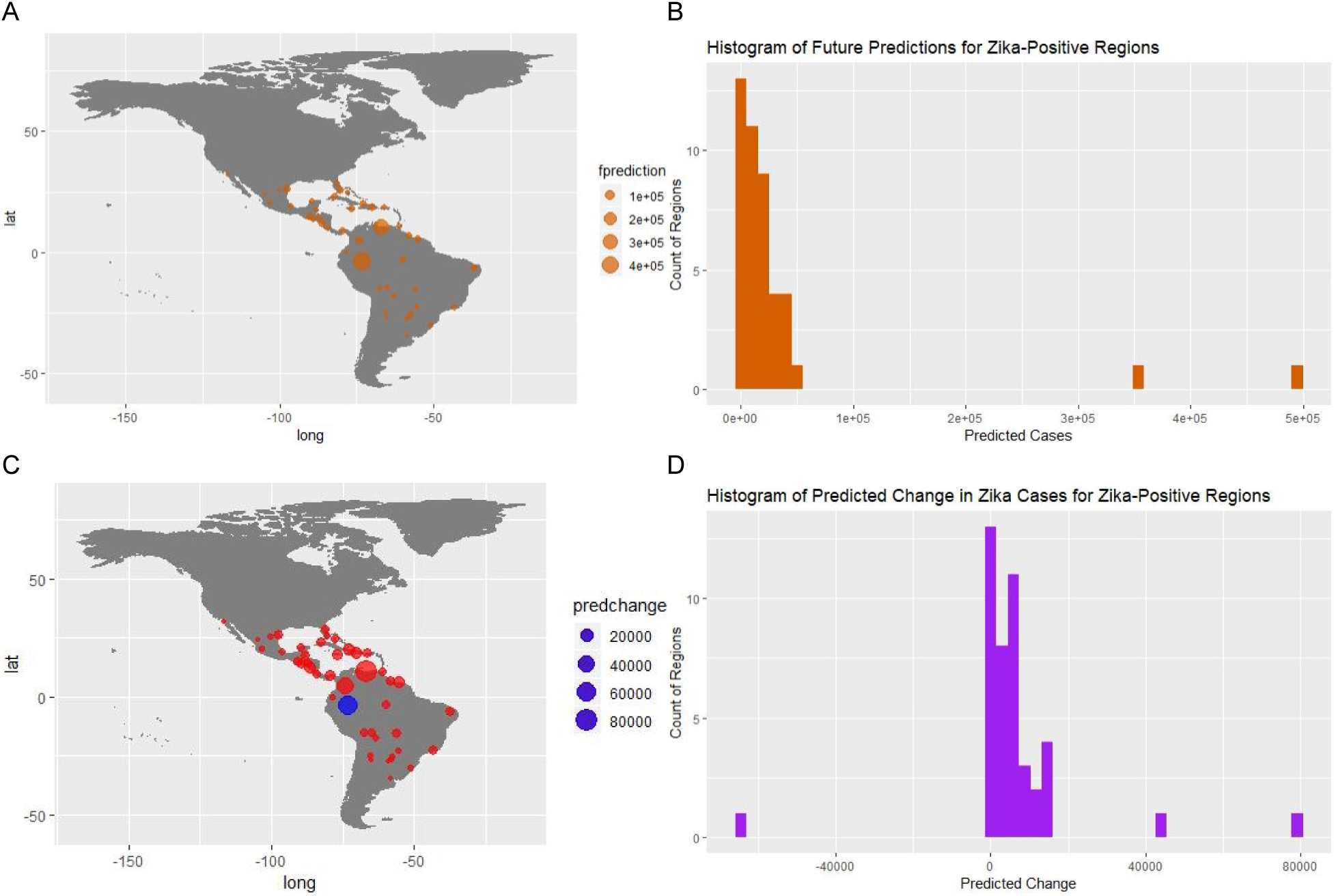
Fig A shows ZINB2’s future predictions for the Zika-positive regions and was produced using the packages maps and ggplot2 in R. In the key, *fprediction* means future prediction. Fig B shows the distribution of future predictions for Zika-positive regions and was produced using ggplot2 in R. Fig C shows ZINB2’s change in predictions for the Zika-positive regions and was produced in the same manner as Fig A. In Fig C, *predchange* means predicted change in cases, and the key shows the absolute value of the predicted change corresponding to the given dot size. On the map, a red dot represents a predicted increase, and a blue dot represents a predicted decrease. Fig D shows the distribution of predicted change in Zika cases for Zika-positive regions and was produced in the same way as Fig B.

## Discussion

In summary, the two top performing models perform well in terms of having acceptable model coefficients, predicting at least one Zika case for all regions that reported Zika, and only predicting at least one case for regions that are habitable for *Ae. aegypti*. Both models were able to predict suitability for *Ae. aegypti* and in turn Zika outbreaks for the chosen subset of regions.

First, we compared the historical averages of the climatic variables used in the model to the values of these variables from 2015-2017, when the Zika outbreak in the Americas occurred. The mean deviations of the 2015-2017 observations from the historical averages were relatively small (between 1 and 2 °C) for the variables *winter.mean*, *summer.mean*, *year.mean*, and *year.diff*. The average yearly mean temperature was higher in the years 2015-2017 than it was in 1980-2005. For *year.min*, the mean deviation from the historical average almost 3 °C, and for *year.max*, the mean deviation was just over 3 °C. On average, *year.min* and *year.mean* were higher and *year.diff* was lower in the years 2015-2017 than in the years 1980-2005, all changes making regions more conducive to Zika outbreaks. None of the selected models used the variable *year.max*, so the deviation in this variable is not concerning. For *year.prec*, the mean deviation was large: almost 1000mm. Most of the places that reported Zika were much drier between 2015 and 2017 than they were between 1980 and 2005, which is unsurprising given the drought reported in Brazil the year than Zika broke out. Since *year.prec* was not used in any of the selected models, the deviation between the climatic average yearly precipitation and the yearly precipitation in 2015-2017 should not impact the performance of the models.

Generally, the ZINB models outperformed the ZIP models. We speculate that the additional variable to deal with over dispersion in the ZINB models contributed to the performance differences. Concerning the performances of the top two models with Zika-free regions, ZINB1 outperformed ZINB2 by only predicting at least one case for regions that either reported the presence of *Ae. aegypti* or were identified as having suitable climates by at least two out of three present potential distributions considered [17][18][21]. ZINB2 predicted just more than one case for Seattle, Washington, which does not have *Ae. aegypti* [33],[62] and is not included in at least two out of three of the likely habitable ranges considered in this paper [17],[18],[21]. Still, this is not too concerning since the prediction is just over one. Both ZINB1 and ZINB2 gave the highest predictions to regions with *Ae. aegypti*. Both ZINB models predicted a median of less than one, although ZINB1 gave more predictions very close to zero, as indicated by its smaller median. ZINB2’s predictions were smaller on average than ZINB1’s, as reflected by their mean predictions.

Concerning the ZINB Models’ performances with regions that did report Zika, both models predicted at least one case for every region. This is crucial, because if a model is to be used to predict where Zika will be a problem in the future, we need to be sure that it does not miss places, resulting in unpreparedness for Zika prevention. For the scaled residuals, ZINB2 had a smaller mean, median, and standard deviation. However, the models’ performances in predicting exact numbers of cases for Zika-positive regions was considered secondarily to their performances in identifying regions that could experience Zika outbreaks. So, we conclude that ZINB1 performed the best.

Looking to the future, the predictions from both the ZINB Models appear to be reasonable: they reaffirm findings that support increased spread of *Ae. aegypti* mosquitoes, and in turn increased climate suitability for Zika outbreaks. Based on the output of these models, the regions that have not yet reported Zika that are most likely to observe an increase in risk for outbreaks include: Los Angeles, California; Louisiana; San Francisco, California; Houston, Texas; Orlando, Florida; Arizona; Uruguay; Cordoba, Argentina; Mississippi; Nevada; and South Carolina.The regions predicted to see the greatest increases in Zika suitability that have already reported Zika include Venezuela; Colombia; Haiti; Nicaragua; the Dominican Republic; Suriname; Panama; Jamaica; Rio de Janeiro, Brazil; Brownsville, Texas; and Puerto Rico.

### Future Considerations

Moving forward, these regions especially should prepare by educating citizens about ways to prevent Zika infections, such as wearing long sleeves, using screens on doors and windows, and using an EPA-registered insect repellant [63]. Economically disadvantaged areas with poor access to water from pipes should work on more secure water storage methods to reduce the creation of *Ae. aegypti* breeding grounds in water containers [64]. Research institutions in these places should record more sightings of *Ae. aegypti* and *Ae. albopictus* and look more into the potential for *Ae. albopictus* to become as much or more of an issue than *Ae. aegypti* in spreading Zika. For example, if the increasing populations of *Ae. albopictus* mosquitoes that have been driving down *Ae. aegypti* populations in the US develop as strong a preference for human blood as *Ae. aegypti* or start to bite primarily inside houses as well, they could become the primary vector for Zika. As these mosquitoes already can survive and reproduce in cooler climates, this could potentially cause more of an issue for Northern regions. In addition, since less is known about the impacts of temperature on *Ae. albopictus*’s feeding and flying habits, more research should be conducted in that area as well.

The BCCCSM1.1 RCP85 climate projections are based on our current trajectory, the highest emissions scenario, which results from not significantly reducing GHGs. These projections suggest that many regions may become more hospitable for *Ae. aegypti*, and many regions that have not seen autochthonous Zika cases before may see them in the future. If actions are taken to reduce GHGs, less severe changes may occur than those projected under this scenario, possibly reducing the risk for Zika to spread to new regions.

In future research, we could use the weather data from the years that the Zika outbreak in the Americas occurred to build the models. Although this data is not available from stations in some of the selected cities in regions that observed Zika (Quito, Ecuador; Georgetown, Guyana; Cayenne, French Guiana; and Port De Paix, Haiti), we could look for the necessary data from other stations in those countries, or we could use historical averages for those regions only. Also, we could increase the temporal resolution of these models’ predictions. Given that monthly or seasonal Zika data is available for enough regions, we might accomplish this by predicting the number of Zika cases per season between the beginning of 2015 and the end of 2017 using the climate data of each season and the season before it [14]. We would also look for socioeconomic data for each region, since socioeconomic factors can contribute to the creation of breeding grounds through water storage methods [64]. We could also see how *Ae. aegypti* density and population density perform as regressors, as they were useful variables in the research of Lo and Park [36]. As a different method of projecting suitability for Zika outbreaks, we could model incidence rate or percentage of population infected, like Huber, Caldwell, and Mordecai had done, rather than modeling cases and including population as one of the regressors [14]. This method may perform better in controlling for differences in population size between regions.

## Abbreviations

| Abbreviation | Term |
| --- | --- |
| Ae. | Aedes |
| BCCCSM1.1 | Beijing Climate Center Climate Systems Model version 1.1 |
| CCAFS | Climate Change, Agriculture, and Food Security |
| CDC | Centers for Disease Control and Prevention |
| ECDC | European Centre for Disease Prevention and Control |
| Fig | Figure |
| GLM | Generalized Linear Model |
| PAHO | Pan American Health Organization |
| RCP8.5 | Representative Concentrations Pathway 8.5 |
| ZINB | Zero-inflated negative binomial |
| ZIP | Zero-inflated Poisson |

## Supporting Information

**S1 Text.** Additional information concerning Generalized Linear Models and Zero-Inflated Models.

**S2 Text.** Performance comparison for all forty models considered, including predictions from best-performing ZIP model.

**S1 Table.** 1980-2005 climate statistics for all 102 regions of interest, with exact latitude and longitude of point used to extract climate data.

**S2 Table.** 2020-2045 climate statistics calculated from bias-corrected BCCCSM1.1 RCP8.5 climate projections.

**S3 Table.** 2015-2017 climate statistics for Zika-positive regions where available. Calculations comparing the 2015-2017 values to the 1980-2005 values.

**S3 Text.** R Codes to: calculate climate statistics from individual data files downloaded from CCAFS for each region, perform the exploratory analysis, impute missing values in Weather Underground data, compare the models considered that were eliminated, compare the two best models, and produce map graphics.

**S1 Zip Folder.** Past and future climate projections and observational data downloaded from CCAFS for each of the 102 regions.

**S2 Zip Folder.** 2015-2017 Weather Underground observations for Zika-positive regions where enough reliable monthly weather observations were available.

## Acknowledgements

SR acknowledges support for the travel from the Pennsylvania State University Eberly College of Science and support for registration for the Society for Mathematical Biology Conference from Lehigh University Department of Mathematics. SR also acknowledges Dr. Ming Li and the Maryland Sea Grant Program funded by NSF grant OCE-1262374 for the opportunity to conduct research in the summer of 2018 that introduced me to climate models and prepared me for this project. SR would also like to individually thank Dr. Teboh-Ewungkem and Dr. Li for their generosity, knowledge, and direction. Dr. Miranda Teboh-Ewungkem acknowledges support via the NSF grant 1815912.

> “We acknowledge the World Climate Research Programme’s Working Group on Coupled Modelling, which is responsible for CMIP, and we thank the climate modeling group, the Beijing Climate Center, for producing and making available their model output. For CMIP the U.S. Department of Energy’s Program for Climate Model Diagnosis and Intercomparison provides coordinating support and led development of software infrastructure in partnership with the Global Organization for Earth System Science Portals.”

## Author Contributions

Conceptualization: Prof. Miranda Teboh-Ewungkem, Samantha Roth

Investigation: Samantha Roth, Prof. Miranda Teboh-Ewungkem

Data Curation: Samantha Roth

Methodology: Samantha Roth

Project Administration: Prof. Miranda Teboh-Ewungkem

Software: Samantha Roth

Supervision: Prof. Miranda Teboh-Ewungkem, Prof. Ming Li

Resources: Prof. Miranda Teboh-Ewungkem

Validation: Samantha Roth, Prof. Miranda Teboh-Ewungkem

Visualization: Samantha Roth

Writing- Original Draft Preparation: Samantha Roth

Writing- Review and Editing: Prof. Miranda Teboh-Ewungkem, Prof. Ming Li, Samantha Roth

